# Morphology and phylogenomics of *Leucomyxa quisqueyensis* sp. nov., a colorless freshwater amoeboid ochrophyte (Stramenopiles) from the Dominican Republic

**DOI:** 10.64898/2026.11.13.766549

**Authors:** Yonas I. Tekle, Saron Ghebrezadik, Angelicia A. Smith, Henry Petit, Cruz Vasquez, Elionor Plasencia, Claudia Reyes, Edian F. Franco

**Affiliations:** Department of Biology, Spelman College, 350 Spelman Lane SW, Atlanta, GA 30314, USA; Department of Basic and Environmental Sciences, Instituto Tecnologico de Santo Domingo (INTEC), Av. Los Proceres 49, Santo Domingo 10602, Dominican Republic; Environmental Genomics Laboratory, Advanced Research Center, Universidad Tecnologica de Santiago (UTESA), Av. Isabel Aguiar 61, Santo Domingo Oeste 21243, Dominican Republic; Department of Research and Innovation, Instituto de Innovacion en Biotecnologia e Industria (IIBI), Santo Domingo 10134, Dominican Republic; Faculty of Sciences, Universidad Autonoma de Santo Domingo (UASD), Av. Alma Mater, Santo Domingo 10105, Dominican Republic

**Author notes:** Correspondence: Yonas I. Tekle.

**Keywords:** Amoeboid stramenopiles, Caribbean freshwater, cryptic plastid, *Leucomyxa*, phylogenomics, reticulopodia

## Abstract

Amoeboid ochrophytes (stramenopiles) exhibit diverse cell forms and plastid states, yet many lineages are known primarily from environmental sequences. We describe *Leucomyxa quisqueyensis* sp. nov., a freshwater amoeboid ochrophyte isolated from a freshwater drainage channel in La Chinola, Santo Domingo Oeste, Dominican Republic. In established cultures, the isolate consistently formed extensive multinucleate reticula, with fine strands linking densely branched regions. The largest networks exceeded 2 mm. Confocal imaging revealed longitudinal microtubule strands along the reticulopodia. Rounded cysts measured 6.47–15.43 µm in diameter. SSU rRNA and nuclear phylogenomic analyses identified *Leucomyxa plasmidifera* as the closest sampled named relative. Their sister relationship received 100% ultrafast bootstrap support in the phylogeny inferred from 124 nuclear protein markers. The two species differed by 2.50% across comparable SSU rRNA positions and by 15.26% across 25,619 comparable amino-acid positions in the nuclear marker alignment. Transcripts with similarity to Tic110 and PPP1, together with matches to plastome-derived protein queries, support the hypothesis that *L. quisqueyensis* retains a cryptic plastid. The new species adds a cultured Caribbean freshwater representative to *Leucomyxa* and provides a basis for comparing reticulopodial organization and plastid evolution among closely related amoeboid ochrophytes.

## Introduction

Stramenopiles encompass photosynthetic algae, phagotrophic protists, and osmotrophic organisms, with forms ranging from unicellular flagellates and amoebae to multicellular seaweeds (Patterson, 1989; Jirsová and Wideman, 2024). Phylogenomic studies have clarified relationships among major stramenopile lineages (Derelle et al., 2016) and resolved the placement of several nonphotosynthetic lineages within the predominantly photosynthetic Ochrophyta (Terpis et al., 2025). Some of these lineages retain reduced plastids after losing photosynthesis, as documented in nonphotosynthetic chrysophytes (Dorrell et al., 2019).

Several ochrophyte lineages include amoeboid forms with branching pseudopodia that can form reticulate networks. The description of the marine amoeboid alga *Synchroma grande*, whose plastid complexes are enclosed by a shared pair of outer membranes, established the class Synchromophyceae (Horn et al., 2007). Its life cycle includes sessile loricate cells, migrating and floating amoebae, and cysts (Koch et al., 2011). Genetically differentiated marine isolates, including isolates from the Caribbean, showed similar morphologies and life histories (Schmidt et al., 2012). The subsequent description of *Guanchochroma wildpretii* expanded the known diversity of amoeboid algae related to *Synchroma* (Schmidt et al., 2015).

The colorless amoeboid strain ATCC PRA-24 was initially studied as *Leukarachnion* sp. because its reticulate networks and cysts resembled those of the freshwater amoeba *Leukarachnion batrachospermi* described by Geitler (1942). Multigene analyses placed the strain within Ochrophyta, close to *Chlamydomyxa* and *Synchroma* (Grant et al., 2009). The relationship of *L. batrachospermi* to ATCC PRA-24 remains unresolved because molecular data are unavailable for *L. batrachospermi*. Barcytė et al. (2024) described ATCC PRA-24 as *Leucomyxa plasmidifera* and documented uninucleate amoebae, multinucleate forms with anastomosing reticulopodia, cysts, and flagellates. They identified a reduced plastid genome and candidate components of the plastid protein import machinery, establishing the presence of a cryptic plastid. Subsequent phylogenomic analyses recovered *Leucomyxa* and *Chlamydomyxa* as sister lineages within an expanded Synchromophyceae that also includes *Synchroma* (Terpis et al., 2025).

The type strain of *Leucomyxa plasmidifera* was isolated from a salt marsh in Virginia, USA. Environmental ITS2 sequences closely matching the type strain were recovered predominantly from soil and soil-associated samples. Environmental small-subunit ribosomal RNA (SSU rRNA) gene sequences affiliated with the broader *Leucomyxa* clade were also reported from compost, lake sediment, and wastewater treatment systems (Barcytė et al., 2024). Much of the known diversity in this clade is represented by environmental sequences, leaving variation in reticulopodial architecture, nuclear organization, and life history poorly characterized.

Here, we characterize an amoeboid ochrophyte isolated from a freshwater drainage channel in La Chinola, Santo Domingo Oeste, Dominican Republic. Using a monoclonal culture, we document reticulopodial architecture and dynamics, nuclear organization, microtubule arrangement, and encystment. We infer the isolate’s phylogenetic placement using SSU rRNA gene sequences and conserved nuclear proteins, and compare its morphology and sequence divergence with those of named relatives to assess its generic placement and species distinction. To investigate possible plastid retention and identify other organelle-related homologs, we screen two transcriptome assemblies against a published protein inventory and a separate set of proteins encoded by the plastid genome, mitochondrial genome, and putative mitochondrial plasmid of *Leucomyxa plasmidifera*.

## Materials and Methods

### Sampling and monoclonal culture

Environmental samples were collected in October 2025 from a freshwater drainage channel located in La Chinola, a densely populated informal settlement adjacent to the Cañada de Guajimía drainage system in Santo Domingo Oeste, Dominican Republic (18.450155° N, 69.991193° W). The sampling site was characterized by substantial anthropogenic disturbance, including visible accumulations of domestic solid waste along the watercourse. Sample aliquots were transferred to plastic Petri dishes containing Deer Park bottled spring water and autoclaved rice grains to promote bacterial growth as a food source for the amoeboid protist. A monoclonal culture was established from a single amoeboid cell isolated from the enrichment. Cultures derived from this isolate were maintained for more than one year for morphological, behavioral, and molecular analyses.

### Light microscopy and immunocytochemistry

Living cells were examined by phase-contrast microscopy. Still images and time-lapse sequences were acquired with a Nikon ECLIPSE Ti2 inverted microscope using NIS-Elements software (Tekle et al., 2026a,b). Filopodial length, reticulopodial branch width, total cell extent, and cyst diameter were measured from phase-contrast images using NIS-Elements. Additional videos were recorded using a smartphone. We documented cell shape, locomotion, pseudopodial extension and branching, formation and reorganization of reticulopodial networks, and encystment.

For whole-mount immunolabeling of microtubules, cells were allowed to adhere to glass chamber slides, rinsed with fresh culture medium, and fixed for 2 min in methanol precooled to −80 °C. The cells were incubated for 1 h with a mouse monoclonal anti-α-tubulin antibody (Life Technologies, 1:500) and then for 45 min with an Alexa Fluor 488-conjugated rabbit anti-mouse IgG (H+L) secondary antibody (Tekle and Wood, 2017). The plasma membrane and nuclear DNA were stained for 5 min with CellMask Orange (Life Technologies, 1:7,000) and DAPI (1:1,000), respectively (Tekle et al., 2014). After rinsing with phosphate-buffered saline, the preparations were mounted under coverslips in Fluoromount-G (Southern Biotech) and imaged with an inverted Zeiss LSM 700 confocal microscope (Carl Zeiss MicroImaging, Göttingen, Germany).

### Transcriptome sequencing and assembly

Cell material was collected by mechanically disrupting branched cells or scraping them from the culture surface. Two preparations were made from the same monoclonal culture, each containing material from approximately 4–20 cells. The preparations were processed separately and sequenced independently as YT-DR-C17 and YT-DR-C18.

cDNA was synthesized and amplified using the SMART-Seq v4 Ultra Low Input RNA Kit (Takara Bio USA). Sequencing libraries were prepared using the Nextera XT DNA Library Preparation Kit (Illumina) following Tekle et al. (2026b). Library DNA concentrations were measured with a Qubit fluorometer using the dsDNA High Sensitivity assay. Libraries were sequenced at Seqcenter (Pittsburgh, PA, USA) in high-output mode, generating paired-end reads of 150 bp.

Read quality and length distributions were assessed with FastQC (Andrews, 2010). Illumina adapters were removed using BBDuk from the BBTools suite (https://github.com/bbushnell/BBTools), and bases with Phred quality scores below 28 were trimmed from both ends of the reads. Reads shorter than 35 bp after trimming were discarded. The retained reads from each preparation were assembled separately *de novo* with rnaSPAdes v4.3.0 (Bushmanova et al., 2019) using default settings, following the read-processing workflow described by Tekle et al. (2020). The assemblies were used to recover SSU rRNA sequences, predict nuclear protein markers, and screen for organelle-associated homologs.

### SSU rRNA phylogenetic analysis

Candidate small-subunit ribosomal RNA (SSU rRNA) gene sequences were identified independently in the YT-DR-C17 and YT-DR-C18 transcriptome assemblies using BLAST (Camacho et al., 2009). The SSU rRNA gene sequence was reconstructed separately from the reads of each run. Reads were recruited if they contained at least two exact 23-nucleotide matches to the corresponding assembly-derived SSU sequence. Local assembly used 51-mers observed at least 20 times and composed entirely of bases with Phred quality scores of at least 20. The C17 reconstruction yielded a 1,844-nt contig that was used as input for the phylogenetic alignment. For deposition (SUB16629855), we selected the 1,716-nt region that could be confidently assigned to the SSU rRNA gene by alignment to the *Leucomyxa plasmidifera* reference sequence (PP766953.1).

The expanded dataset included 59 taxa, including environmental sequences, and was used to assess the isolate’s placement within the broader *Leucomyxa* clade. The 23-taxon dataset was a subset of the expanded dataset and was used to assess relationships between the isolate and named taxa. Both datasets included the SSU rRNA gene sequence of *Leucomyxa plasmidifera* ATCC PRA-24 (PP766953.1) (Barcytė et al., 2024).

Sequences in the expanded dataset were aligned with MAFFT v7.526 using L-INS-i (Katoh and Standley, 2013). The alignment was trimmed to the span of PP766953.1, and columns with unambiguous nucleotides in fewer than 50% of the sequences were excluded. The 23-taxon subset was extracted from the filtered alignment without further column filtering, so both datasets retained the same 1,679 positions.

Maximum-likelihood trees were inferred with IQ-TREE v2.4.0 (Minh et al., 2020). ModelFinder selected TIM3e+I+R3 for the 59-taxon dataset using the Bayesian information criterion (Kalyaanamoorthy et al., 2017), and the same model was used for the 23-taxon analysis. Branch support was assessed with 1,000 ultrafast bootstrap replicates (Hoang et al., 2018) and 1,000 SH-aLRT replicates (Guindon et al., 2010). Both trees were rooted with *Nannochloropsis oceanica*.

Uncorrected pairwise divergence was calculated as the number of mismatches divided by the number of aligned positions with unambiguous nucleotides in both sequences and expressed as a percentage. For the common-site divergence comparisons, we retained the 1,531 alignment positions with unambiguous nucleotides in all five focal sequences: the La Chinola isolate, *Leucomyxa plasmidifera*, *Synchroma pusillum*, *Chlamydomyxa labyrinthuloides*, and *Synchroma grande* (Table S1).

### Nuclear phylogenomics and protein divergence

We predicted open reading frames separately from the YT-DR-C17 and YT-DR-C18 transcriptome assemblies, which represent two sequencing runs of the same La Chinola isolate. We used TransDecoder v6.0.0 (Haas, 2026) with a minimum open reading frame length of 50 amino acids and with start-site refinement disabled. We retained the highest-ranked prediction for each transcript. We assessed conserved eukaryotic marker recovery in each predicted protein set using BUSCO v6.1.0 in protein mode with the eukaryota_odb10 dataset (Manni et al., 2021). C18 supplied the isolate’s protein sequences for the final phylogenomic and divergence analyses.

We used BLASTP (Camacho et al., 2009) to search the predicted C17 and C18 proteins against 240 reference ortholog families from Terpis et al. (2025). We obtained the ortholog and paralog reference sequences from the `Stram_PF_release.tar.gz` archive in the associated Dryad dataset (doi:10.5061/dryad.1g1jwsv5v). For each query, the top-scoring ortholog hit was required to span at least 100 amino acids, cover at least 60% of the query and 30% of the reference sequence, and have an E-value ≤ 10□¹□. We excluded a query if its best hit to another ortholog family had a bit score at least 90% of the bit score of its top ortholog hit. We excluded a family from the C18 marker set if its second-ranked eligible candidate had a bit score at least 90% of that of the highest-ranked candidate. We then screened the remaining C18 candidates against the pooled reference paralog sequences and excluded candidates whose best paralog hit had a bit score at least 95% of that of their top ortholog hit.

Each eligible orthologous marker family was aligned using MAFFT v7.526 with the `--auto` option (Katoh and Standley, 2013). Each alignment contained at least 25 sequences, including a C18 protein and representatives of all three focal reference strains: *Leucomyxa plasmidifera* ATCC PRA-24, *Chlamydomyxa montana* SAG 36.96, and *Synchroma pusillum* CCMP3072. Alignment columns were retained if they contained an amino acid residue in C18 and had at least 70% occupancy among reference sequences, with gaps and ambiguous residues (X) treated as missing data. Markers with at least 100 retained alignment positions were concatenated into a supermatrix, with missing loci represented by gaps. Reference taxa with less than 35% occupancy in the concatenated alignment were excluded, except for the three focal reference strains. The final concatenated dataset comprised 124 protein markers, 31,040 aligned amino acid positions, and 38 taxa (Table S2).

For each retained marker with a corresponding C17 candidate, we locally aligned the C17 and C18 predictions using BLOSUM62 with gap-opening and gap-extension scores of −10 and −0.5, respectively. We excluded positions with a gap or an X in either sequence from the percent identity calculation and the count of comparable positions. For overlaps with at least 50 comparable amino-acid positions, we recorded whether the predictions were identical at all of those positions (Table S2A).

We inferred a maximum-likelihood tree with IQ-TREE v3.0.1 (Wong et al., 2026) using a single LG+G4 model across the concatenated alignment (Le and Gascuel, 2008). We assessed branch support with 1,000 ultrafast bootstrap replicates (Hoang et al., 2018) and 1,000 SH-like approximate likelihood-ratio test replicates (Guindon et al., 2010). We rooted the tree for display on the branch leading to the four sampled Bigyra taxa (Fig. 4). For each individual marker tree, we excluded sequences with fewer than 30 amino-acid residues after disregarding gaps and X, then used IQ-TREE’s fast maximum-likelihood search under LG+G4. We counted the marker trees that recovered the La Chinola isolate and *L. plasmidifera* ATCC PRA-24 as an exclusive pair (Table S2C).

We calculated uncorrected pairwise amino-acid divergence among the La Chinola isolate and the three focal references. All six comparisons used the same 25,619 alignment positions, each containing one of the 20 standard amino acids in all four taxa. For each pair, divergence was the percentage of these positions at which the two sequences differed (Table S3).

### Screening for plastid and mitochondrial homologs

To identify candidate homologs of proteins associated with plastid function and metabolism, we screened the YT-DR-C17 and YT-DR-C18 transcriptome assemblies separately using TBLASTN (Camacho et al., 2009) with an E-value cutoff of 10 □□. The 94 protein queries were drawn from the published inventory for *Leucomyxa plasmidifera* ATCC PRA-24 (table S5 of Barcytė et al., 2024) and included proteins associated with plastid protein import, transcription, translation, and metabolism.

In a separate screen, we searched both assemblies with 125 protein queries translated from annotated coding sequences in three GenBank records for the same strain. The query set comprised 47 proteins from the plastid genome (PP760456.1), 49 from the mitochondrial genome (PP760457.1), and 29 from the putative mitochondrial plasmid (PP760458.1). For both screens, we recorded whether each reference query yielded at least one hit in each assembly (Tables S4 and S5, respectively).

To extend the phenylalanine hydroxylase (PAH) candidate, we searched the oriented C17 and C18 contigs for an exact suffix–prefix overlap of at least 100 nt and joined the contigs across the shared sequence. We examined conserved domains in selected candidates using `hmmscan` in HMMER v3.4 (Eddy, 2011) and applied the family-specific gathering thresholds of the Pfam profiles retrieved through InterPro (Paysan-Lafosse et al., 2025).

For the expanded gene-family phylogenies, we aligned candidate proteins and reference homologs with MAFFT v7.526 using `--auto`. We retained columns with a residue other than a gap or X in the focal sequence and at least 50% occupancy among the reference sequences. After trimming, we excluded sequences with fewer than 30 residues or less than 35% occupancy, treating gaps and X as missing. We inferred maximum-likelihood trees with IQ-TREE v3.0.1 under LG+G4 and assessed branch support with 1,000 ultrafast bootstrap replicates and 1,000 SH-aLRT replicates. Reference comparisons included homologs from other ochrophytes, including taxa sampled by Terpis et al. (2025). Where relevant, we also compared candidates with bacterial and fungal homologs to evaluate possible culture-associated origins. We classified sequence matches as candidate homologs and treated protein localization and organelle retention as hypotheses requiring independent evidence.

## Results

### Morphology and development in culture

The La Chinola isolate formed networks of fine, anastomosing reticulopodia. Smaller forms bore tapering projections and had only a few spaces between branches (Fig. 1A–C). Larger networks comprised densely interconnected regions linked by long, slender strands (Fig. 1D–E). The spaces enclosed by intersecting branches ranged from small and rounded to large and irregular or polygonal. The largest networks extended more than 2 mm (Video S1). Fine peripheral filopodia were difficult to distinguish against the bacterial background under phase contrast and measured 1–5 µm in length (mean 3.7 µm, *n* = 51). The reticulopodial branches were 1–12 µm wide (mean 5 µm, *n* = 34). Time-lapse recordings showed branches extending and retracting as the network moved across the substrate, continually reshaping the spaces between them (Video S1).

**Fig. 1.**
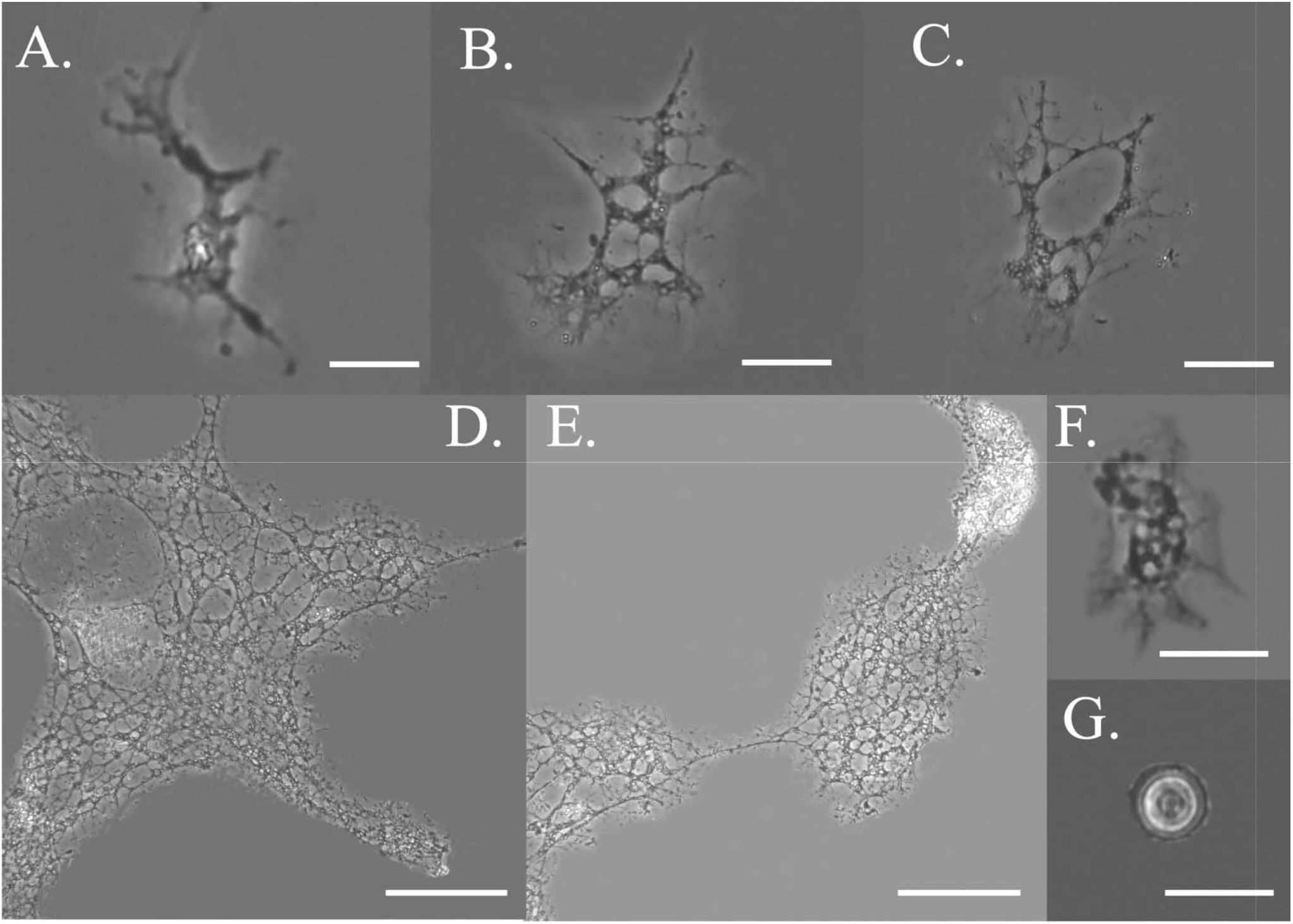
Vegetative morphology and cyst of *Leucomyxa quisqueyensis* sp. nov. in culture. Phase-contrast micrographs show (A–C) smaller reticulate forms with tapering projections, (D–E) extensive reticula with densely branched regions connected by slender strands, (F) a small amoeboid cell with a discernible cell body and peripheral projections, and (G) a rounded cyst with two discernible concentric wall boundaries. Scale bars are 20 µm in A–C and F–G and 100 µm in D–E.

Reticulopodia fragmented readily when disturbed, including during scraping for subculture. Some detached fragments gave rise to small uninucleate amoeboid cells. Shortly after subculture, cells with one or a few nuclei had a discernible cell body and commonly spanned approximately 30–117 µm in total extent, including their projections (Fig. 1F). Extensive multinucleate networks predominated in established cultures. DAPI-labeled nuclei lay alongside α-tubulin-labeled (microtubules) strands in narrow branches and were distributed throughout broader regions of the reticulum (Fig. 2). Approximately 30 nuclei were distinguishable in the network shown in Fig. S1A–B. Nuclear diameter ranged from 0.98 to 2.42 µm (mean 1.65 µm, *n* = 61).

**Fig. 2.**
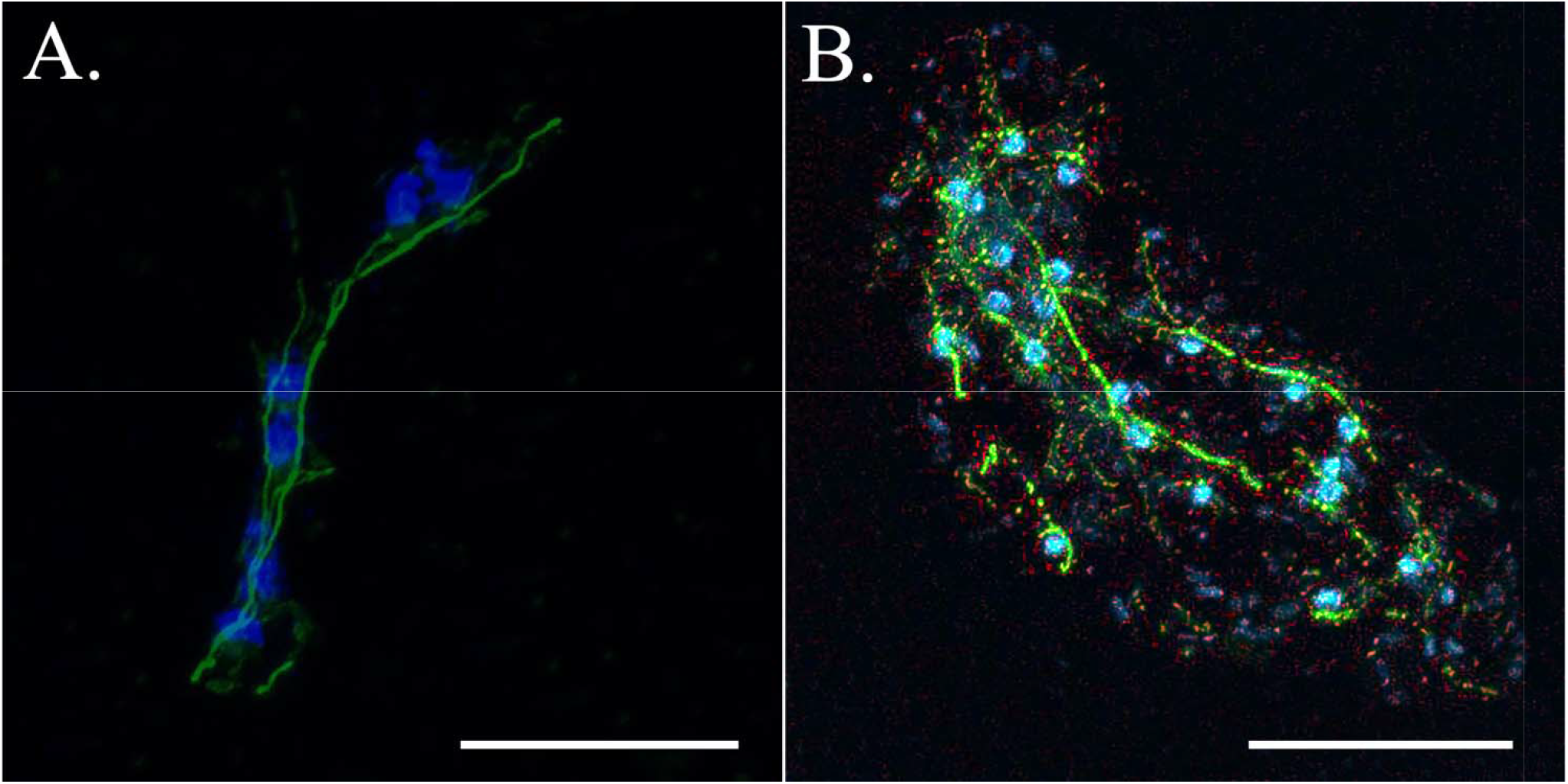
Nuclear distribution and microtubule organization in vegetative *Leucomyxa quisqueyensis* sp. nov. Confocal micrographs show (A) DAPI-labeled nuclei along a narrow branch with longitudinal α-tubulin labeling and (B) nuclei distributed among α-tubulin-labeled strands in a broader region of the reticulum. α-Tubulin immunolabeling is shown in green, plasma membrane staining in red where displayed, and DAPI staining in blue. Scale bars are 20 µm.

Anti-α-tubulin immunolabeling of microtubules revealed longitudinal strands along the reticulopodial branches (Fig. 2 and Fig. S1A). The apparent thickness of the strands was consistent with microtubule bundles. Individual microtubules could not be resolved in the confocal images.

### Cysts and observations of flagellates

Reticulopodial networks were most extensive in cultures with limited bacterial overgrowth. Cysts occurred in older cultures with abundant bacteria and accumulated debris. They were rounded, with two discernible concentric wall boundaries, and measured 6.47–15.43 µm in diameter (mean 11.44 µm, *n* = 64). A representative cyst is shown in Fig. 1G. Confocal imaging revealed two DAPI-positive bodies consistent with nuclei in the upper cyst and at least three in the lower cyst (Fig. S1C–D).

Flagellates appeared once during a year of culture observations. Their relationship to the La Chinola isolate remained uncertain because flagellates had also been present in the original mixed culture.

### SSU rRNA phylogenetic placement and sequence divergence

In the 23-taxon SSU rRNA analysis, the La Chinola isolate was recovered as sister to *Leucomyxa plasmidifera* (PP766953.1), with 99.3% SH-aLRT and 99% ultrafast bootstrap support (Fig. 3). The *Leucomyxa* lineage grouped with representatives of Chlamydomyxales and Synchromales, consistent with the SSU rRNA phylogeny of Barcytė et al. (2024). In the expanded 59-taxon analysis, the isolate and *L. plasmidifera* fell within a broader clade that also contained environmental sequences. Support for this clade was limited (30.9% SH-aLRT and 66% ultrafast bootstrap support, Fig. S2). Within this clade, the isolate formed a distinct branch sister to a subclade containing *L. plasmidifera* and several environmental sequences.

**Fig. 3.**
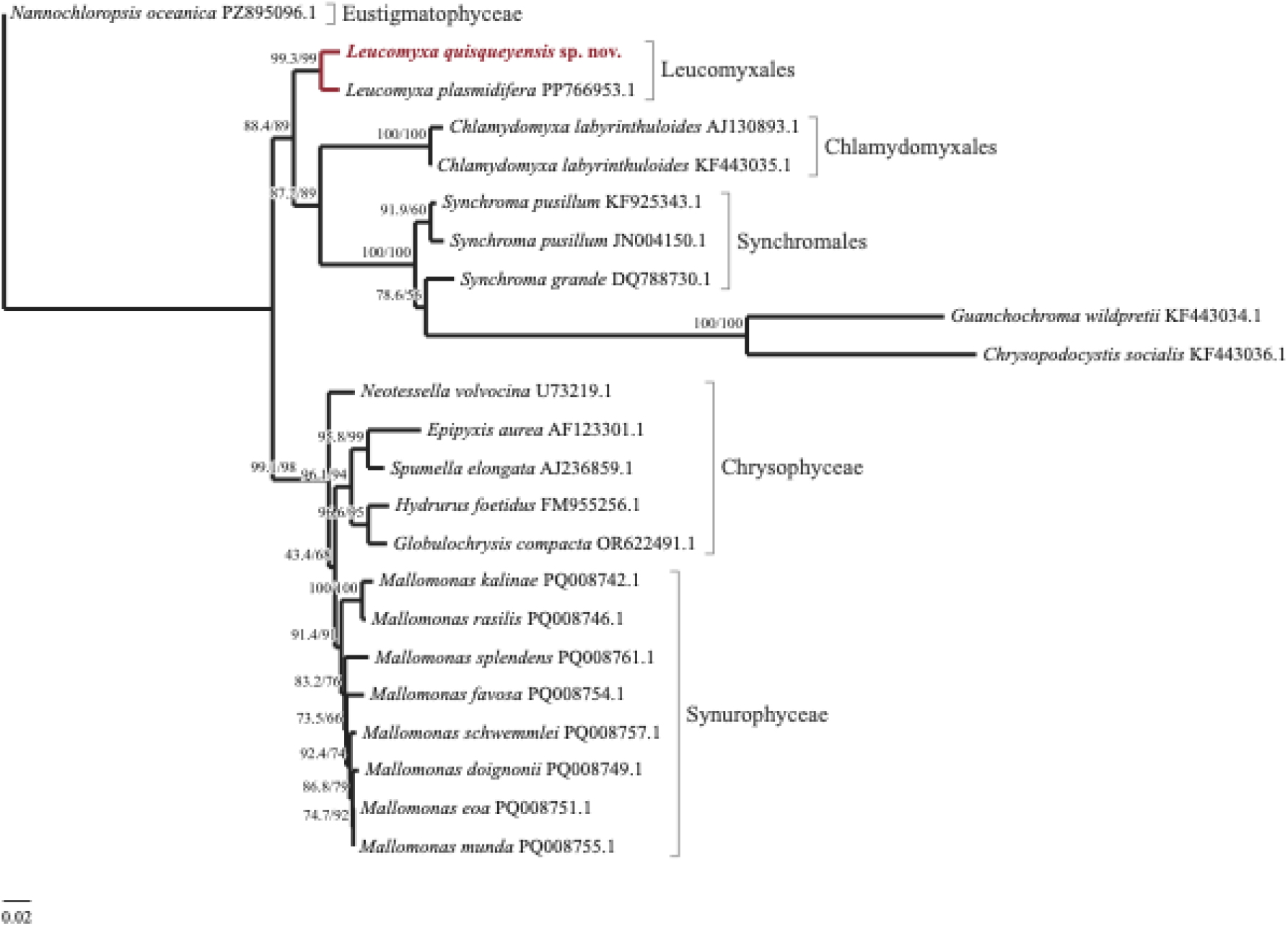
SSU rRNA phylogenetic placement of *Leucomyxa quisqueyensis* sp. nov. among named ochrophytes. The maximum-likelihood tree was inferred from 23 sequences and 1,679 aligned nucleotide positions and rooted for display with *Nannochloropsis oceanica*. Branch labels show SH-aLRT and ultrafast bootstrap support, in that order. The stem branch of the clade comprising *Guanchochroma wildpretii* and *Chrysopodocystis socialis* is drawn at half its inferred length for display. The scale bar represents 0.02 substitutions per nucleotide site and applies to all other branches.

The isolate differed from *L. plasmidifera* at 42 of 1,678 pairwise comparable nucleotide positions (2.50% uncorrected divergence). Across the 1,531 positions comparable among all five focal sequences, the isolate differed from *L. plasmidifera* at 26 positions (1.70%). On the same set of sites, divergence from *Synchroma pusillum*, *Chlamydomyxa labyrinthuloides*, and *Synchroma grande* was 7.64%, 7.97%, and 8.36%, respectively (Table S1).

### Nuclear phylogenomic placement and amino-acid divergence

BUSCO identified 44 complete markers in the C18 predicted protein set and eight in C17. C18 supplied the isolate’s protein sequences for nuclear phylogenomics and amino-acid divergence, while C17 supplied its SSU rRNA sequence for phylogenetic inference. Both assemblies were used in the organelle-related protein screens.

The maximum-likelihood analysis of 124 nuclear protein markers recovered the La Chinola isolate as sister to *Leucomyxa plasmidifera*, with 100% SH-aLRT and 100% ultrafast bootstrap support (Fig. 4). *Chlamydomyxa montana* SAG 36.96 was sister to this pair, and *Synchroma pusillum* CCMP3072 was sister to the resulting three-taxon clade. Each of these two additional relationships also received 100% SH-aLRT and 100% ultrafast bootstrap support. The isolate and *L. plasmidifera* formed an exclusive pair in 107 of the 124 individual maximum-likelihood marker trees (Table S2C).

**Fig. 4.**
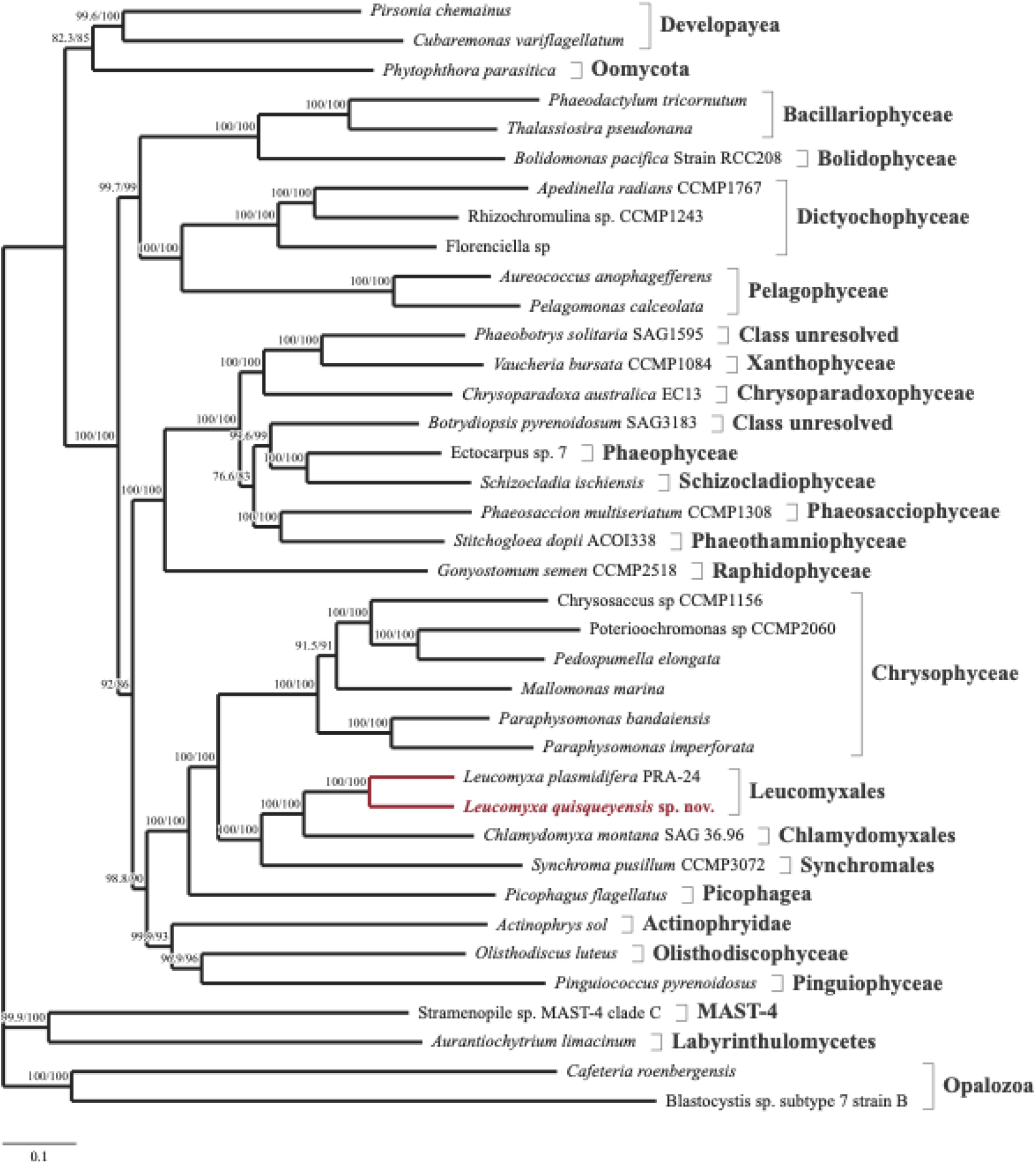
Nuclear phylogenomic placement of *Leucomyxa quisqueyensis* sp. nov. The maximum-likelihood tree was inferred under the LG+G4 model from 124 concatenated nuclear protein markers comprising 31,040 aligned amino-acid positions across 38 taxa. The tree was rooted for display on the branch leading to the four sampled Bigyra taxa. Branch labels show SH-aLRT and ultrafast bootstrap support, in that order, based on 1,000 replicates each. The scale bar represents 0.1 substitutions per amino-acid site.

The overall topology including clade comprising the sampled diatoms and bolidophyte was sister to the clade comprising the sampled dictyochophytes and pelagophytes, consistent with the grouping of these lineages in Diatomista in previous phylogenomic studies (Derelle et al., 2016, Cho et al., 2024).

Across the 25,619 amino-acid positions comparable among all four focal taxa, the isolate differed from *L. plasmidifera* at 3,909 positions (15.26% uncorrected divergence). On the same set of sites, it differed from *C. montana* at 5,983 positions (23.35%) and from *S. pusillum* at 7,506 positions (29.30%) (Table S3). To assess sequence agreement between the two transcriptome assemblies, we compared the retained C18 markers with their C17 counterparts. Of the 124 markers, 82 had at least 50 comparable amino-acid positions. For 56 of those 82 markers, the C17 and C18 predictions were identical at every comparable position (Table S2A).

### Recovery of plastid- and organelle-related protein candidates

Screening the two transcriptome assemblies against 94 published *Leucomyxa plasmidifera* protein queries yielded candidate matches to 74 queries in C17 and 72 in C18. Of the 94 queries, 67 yielded matches in both assemblies, seven only in C17, and five only in C18. Overall, 79 queries yielded a candidate in at least one assembly, and 15 yielded no match in either assembly (Table S4).

C18 yielded partial matches to two proteins associated with plastid protein import. The Tic110 candidate shared 42.1% amino-acid identity with the reference over 140 residues, covering approximately 10% of the query. The PPP1 candidate shared 80.6% identity over 67 residues, covering approximately 22% of the query (Table S4).

The oriented C17 and C18 contigs matching phenylalanine hydroxylase (PAH) shared an exact 382-nucleotide overlap. Joining the contigs at this overlap yielded an inferred 1,647-nucleotide sequence containing a putative open reading frame encoding 416 amino acids. The predicted protein contained a biopterin-dependent hydroxylase domain and the conserved iron-binding residues H255, H260, and E300, supporting its assignment to the PAH family (Fig. S3A–B).

Three separate C18 contigs matched distinct regions of the published 1,004-amino-acid PAPSS reference protein. One matched the APS kinase and PUA regions, another matched the ATP sulfurylase region, and the third matched the pyrophosphatase region. Whether these regions occur within a single transcript in the isolate remains unresolved (Fig. S3C).

In a separate screen of 125 proteins translated from *L. plasmidifera* reference coding sequences, matches were recovered for 29 of the 47 plastid genome queries in C17 and for 23 in C18. Of the 49 mitochondrial genome queries, 39 yielded matches in each assembly. Of the 29 queries from the putative mitochondrial plasmid, seven yielded matches in C17 and eight in C18 (Table S5). The genomic origin and subcellular localization of these candidates remain unresolved, as does the presence of a plastid or mitochondrial plasmid in the La Chinola isolate.

## Discussion

### Phylogenetic placement and species distinction

The SSU rRNA and nuclear protein analyses identify *Leucomyxa plasmidifera* as the closest sampled named relative of the La Chinola isolate. In the nuclear tree, the two isolates formed a clade, with *Chlamydomyxa montana* and *Synchroma pusillum* as successive sister lineages, consistent with the relationships recovered by Terpis et al. (2025). With environmental sequences included, the La Chinola isolate formed a distinct branch within the broader *Leucomyxa* assemblage. Limited support for this assemblage leaves the isolate’s relationships to individual environmental lineages unresolved. Barcytė et al. (2024) documented substantial SSU diversity in the assemblage. The La Chinola culture links a branch of that diversity to observations of living cells, developmental stages, and nuclear protein sequences.

The La Chinola isolate differed from *L. plasmidifera* by 2.50% across pairwise comparable SSU rRNA sites and by 15.26% across the amino-acid positions shared by all four focal taxa in the nuclear marker panel. The sequence differentiation in both datasets, together with the larger cysts and recurrent development of extensive reticula observed in the La Chinola culture, supports recognition of the isolate as a distinct species of *Leucomyxa*. Additional isolates would help determine how consistently these morphological characters distinguish the species.

### Reticulopodial development and cytoskeletal organization

The La Chinola isolate and *L. plasmidifera* share anastomosing reticulopodia and multinucleate vegetative forms. Grant et al. (2009) reported a large network in the strain later described as *L. plasmidifera*. Barcytė et al. (2024) observed smaller, short-lived networks in the same strain and suggested that culture conditions could explain the difference. In established La Chinola cultures, extensive reticula formed consistently. Delicate strands connected densely branched regions, and the largest networks exceeded 2 mm. The regular development of this architecture is a potentially useful distinguishing character, although its diagnostic value would be best assessed under matched culture conditions. Cyst diameter provides a more direct comparison. Cysts of the La Chinola isolate measured 6.47–15.43 µm in diameter (mean 11.44 µm, *n* = 64), with only a narrow overlap with the published range of 2.5–7.0 µm for *L. plasmidifera* (Barcytė et al., 2024).

Within the extensive La Chinola reticulum, α-tubulin labeling was concentrated in longitudinal strands whose apparent thickness was consistent with microtubule bundles. A comparable arrangement occurs in the branching amoebozoan *Acramoeba dendroida*, where bundles run along broader branches and individual microtubules extend into fine projections (Tekle and Williams, 2016). Other amoebozoans illustrate how differently microtubules can be organized around the nucleus. *Stereomyxa ramosa*, another branching amoeba, has a prominent perinuclear microtubule-organizing center (MTOC) and individual or looping microtubules in its projections (Tekle and Wood, 2017). In *Stratorugosa tubuloviscum*, microtubules associated with a perinuclear MTOC can wrap around the nucleus (Melton et al., 2019). The nucleus-associated acentriolar centrosome of *Dictyostelium discoideum* provides another example of an organized cytoplasmic array (Gräf et al., 2021). These comparisons distinguish the arrangement of microtubules within projections from the location of their nucleation. The La Chinola images establish the arrangement of labeled microtubule strands along branches. The sites of microtubule nucleation and the presence of a discrete MTOC remain unresolved.

The occurrence of interconnected projections in ochrophytes, variosean amoebozoans, and foraminifera suggests convergence in cell architecture across distantly related eukaryotes (Grant et al., 2009, Berney et al., 2015, Travis and Bowser, 1988). In foraminifera, reticulopodial microtubules serve as tracks for rapid bidirectional organelle transport (Travis and Bowser, 1988). In the La Chinola isolate, nuclei occurred alongside microtubule strands in narrow branches. This spatial association raises the question of how nuclei are redistributed as a millimetre-scale reticulum extends, retracts, and fragments. The La Chinola isolate thus provides an ochrophyte model for testing whether independently evolved amoeboid architectures rely on similar mechanisms of cytoskeletal organization and intracellular transport.

### Cryptic plastid retention and organelle evolution

The close relationship to *L. plasmidifera*, which retains a reduced plastome, makes inheritance of a nonphotosynthetic plastid a plausible hypothesis for the La Chinola lineage (Barcytė et al., 2024). The Tic110- and PPP1-like transcripts add evidence from the isolate itself. Tic110 is associated with protein import at the inner plastid envelope. PPP1 participates in protein import into the apicoplast of *Toxoplasma gondii*, providing a functional comparison across red-alga-derived complex plastids. Its role in ochrophytes remains unresolved (Sheiner et al., 2011, Barcytė et al., 2024). These candidates are informative because nonphotosynthetic complex plastids can depend on continued import of nucleus-encoded proteins to maintain their remaining functions (Maier et al., 2015, Sheiner et al., 2011, Barcytė et al., 2024).

Both low-input transcriptomes recovered incomplete sets of conserved markers. In this context, the positive matches to import-associated proteins are more informative than individual nondetections, although their short alignments leave full protein architecture and targeting unresolved. The separate screen recovered matches to proteins encoded by the *L. plasmidifera* plastome. The genomic origin of the matching transcripts in the La Chinola isolate remains unresolved. The inferred PAH extension and PAPSS-related fragments support comparisons of individual metabolic proteins between the isolate and *L. plasmidifera* (Fig. S3). Their roles in plastid metabolism remain to be established. Taken together, phylogenetic proximity to a plastid-bearing species, recovery of import-associated transcripts, and matches to plastome-derived queries support cryptic plastid retention as a working hypothesis. Localization of the candidate proteins to a plastid or recovery of an isolate-associated plastome would strengthen that inference.

Closely related lineages can provide a comparison of what persists after photosynthesis is lost. Nonphotosynthetic *Spumella* retains a reduced plastome, whereas a plastid is inferred to persist after plastome loss in *Paraphysomonas* (Dorrell et al., 2019). Terpis et al. (2025) inferred independent losses of the entire plastid in *Actinophrys sol* and *Picophagus flagellatus*. Outside ochrophytes, cryptic plastids in *Rhodelphis* and *Euglena longa* illustrate further variation in organelle reduction and retention (Gawryluk et al., 2019, Füssy et al., 2020). These cases show that photosynthesis, plastid DNA, and the plastid itself can be lost separately. Determining the plastid status of the La Chinola isolate would help establish whether plastid retention extends across these two sampled lineages and, if so, which plastid functions they share.

The mitochondrial-plasmid matches raise a separate question about the evolution of extrachromosomal DNA in *Leucomyxa*. Fungal mitochondrial plasmids provide a precedent for exchange between mitochondrial genomes and plasmids, including transfer of mitochondrial tRNA genes to plasmids (Nieuwenhuis et al., 2023). The transcript matches recovered here identify candidates for comparison with the putative *L. plasmidifera* plasmid (Barcytė et al., 2024).

Establishing whether the corresponding sequences occur on a discrete extrachromosomal molecule would clarify whether the two lineages share a mitochondrial plasmid and, if so, how their plasmid gene content differs.

### Caribbean freshwater diversity and persistence in culture

Cultured amoeboid ochrophytes provide a way to compare cell architecture and organelle biology within a phylogenetic framework. *Synchroma grande* and *S. pusillum* possess conspicuous plastid complexes, and *L. plasmidifera* retains a reduced, nonphotosynthetic plastid (Horn et al., 2007, Schmidt et al., 2012, Barcytė et al., 2024). The La Chinola isolate adds a freshwater representative for which live reticulopodial behavior, nuclear organization, and encystment can be examined alongside molecular data. Such observations give biological context to the environmental sequences that account for much of the diversity currently known near *Leucomyxa*.

The two isolates were recovered from different habitats. The *L. plasmidifera* reference strain originated in a Virginia salt marsh and has been maintained in seawater-based medium (Grant et al., 2009, Barcytė et al., 2024). Closely matching ITS2 signatures were recovered predominantly from soils and soil-associated samples (Barcytė et al., 2024). The La Chinola isolate was recovered from a freshwater drainage channel in La Chinola, Santo Domingo Oeste. In a preliminary culture trial, direct transfer of the La Chinola isolate to 3% artificial seawater was followed by network disruption and cell death (data not shown). Together with its freshwater origin and sustained growth in freshwater culture, this response is consistent with freshwater adaptation. The salinity tolerance ranges of the two isolates remain to be determined.

The recovery of this isolate from La Chinola documents its occurrence in a Caribbean urban stream where household waste was visible at the collection site. Urban streams can experience concurrent changes in hydrology, water chemistry, and biological communities (Walsh et al., 2005). In culture, some reticulopodial fragments gave rise to small amoeboid cells after mechanical disturbance, and cysts formed in older cultures with abundant bacteria and accumulated debris. These observations document recovery after fragmentation and the formation of a potential resting stage. Their contributions to survival under specific environmental stresses remain to be tested. Sampling Caribbean urban and less disturbed freshwaters would help establish the distribution of *Leucomyxa*-like lineages and link environmental detections to traits that can be measured in culture.

## Taxonomic treatment

### *Leucomyxa quisqueyensis* Tekle and Franco, sp. nov

#### Differential diagnosis

Freshwater amoeboid species of *Leucomyxa*. In established cultures, it consistently forms extensive multinucleate reticula in which delicate, anastomosing branches connect densely branched regions. The largest networks exceed 2 mm (Video S1). Rounded cysts with two discernible wall boundaries measure 6.47–15.43 µm in diameter (mean 11.44 µm, *n* = 64), generally larger than those reported for *L. plasmidifera* (2.5–7.0 µm, Barcytė et al., 2024). The isolate differs from *L. plasmidifera* at 42 of 1,678 pairwise comparable SSU rRNA positions (2.50%) and at 3,909 of 25,619 amino-acid positions (15.26%) shared by the four focal taxa in the nuclear protein alignment (Tables S1 and S3).

#### Description

Shortly after subculture, cells with one or a few nuclei have a discernible cell body and typically span approximately 30–117 µm, including their projections. In established cultures, extensive multinucleate reticula predominate. Peripheral filopodia measure 1–5 µm in length (mean 3.7 µm, *n* = 51), and reticulopodial branches measure 1–12 µm in width (mean 5 µm, *n* = 34). Nuclear diameter is 0.98–2.42 µm (mean 1.63 µm, *n* = 61). Longitudinal strands labeled with anti-α-tubulin follow the reticulopodial branches. Cysts occur in older cultures. A flagellated stage remains unconfirmed (Figs. 1–2 and S1).

#### Type locality

Freshwater drainage channel in La Chinola, a densely populated informal settlement adjacent to the Cañada de Guajimía drainage system in Santo Domingo Oeste, Dominican Republic (18.450155° N, 69.991193° W). The sample was collected in October 2025.

#### Holotype

Cryopreserved material from a monoclonal culture established from a single amoeboid cell isolated from the type locality. The material is held in cryogenic storage in the Tekle Laboratory Culture Collection, Department of Biology, Spelman College, Atlanta, Georgia, USA.

#### Photographic and video documentation

Phase-contrast images document vegetative forms and cyst morphology (Fig. 1), confocal images document nuclear and microtubule organization (Figs. 2 and S1), and time-lapse recordings document reticulopodial dynamics (Video S1).

#### Genetic reference material

A 1,716-nt SSU rRNA gene sequence reconstructed from the C17 sequencing run of the type isolate serves as the molecular reference for the species (GenBank accession SUB16629855).

#### Etymology

The epithet *quisqueyensis* sp. nov. is a Latinized geographic adjective derived from Quisqueya, a name used for the Dominican homeland.

## Acknowledgments

This work was supported by the Simons Foundation Fellow Award (SFA-23-5) and the National Science Foundation Excellence in Research (EiR) Award (2401946), both to Yonas I. Tekle, and by the Ministerio de Educación Superior, Ciencia y Tecnología (MESCyT) of the Dominican Republic through the Fondo Nacional de Innovación y Desarrollo Científico y Tecnológico (FONDOCYT, Grant No. 2024-2-2B3-0919 to Edian F. Franco). We thank Christon J. M. Racoma and Kierstyn Landry for their assistance with culturing the isolate and the Executive Directorate of the Instituto de Innovación en Biotecnología e Industria (IIBI) for its institutional, administrative, and logistical support throughout the project.

## Data Availability Statement

Raw RNA-seq reads from the two sequencing runs of the *Leucomyxa quisqueyensis* type isolate are available in the NCBI Sequence Read Archive under accessions [PRJNA1560882 for YT-DR-C17] and [PRJNA1560882 for YT-DR-C18]. The 1,716-nt SSU rRNA gene sequence reconstructed from YT-DR-C17 is available in GenBank under accession [SUB16629855].

## Supplementary figures

**Fig. S1.**
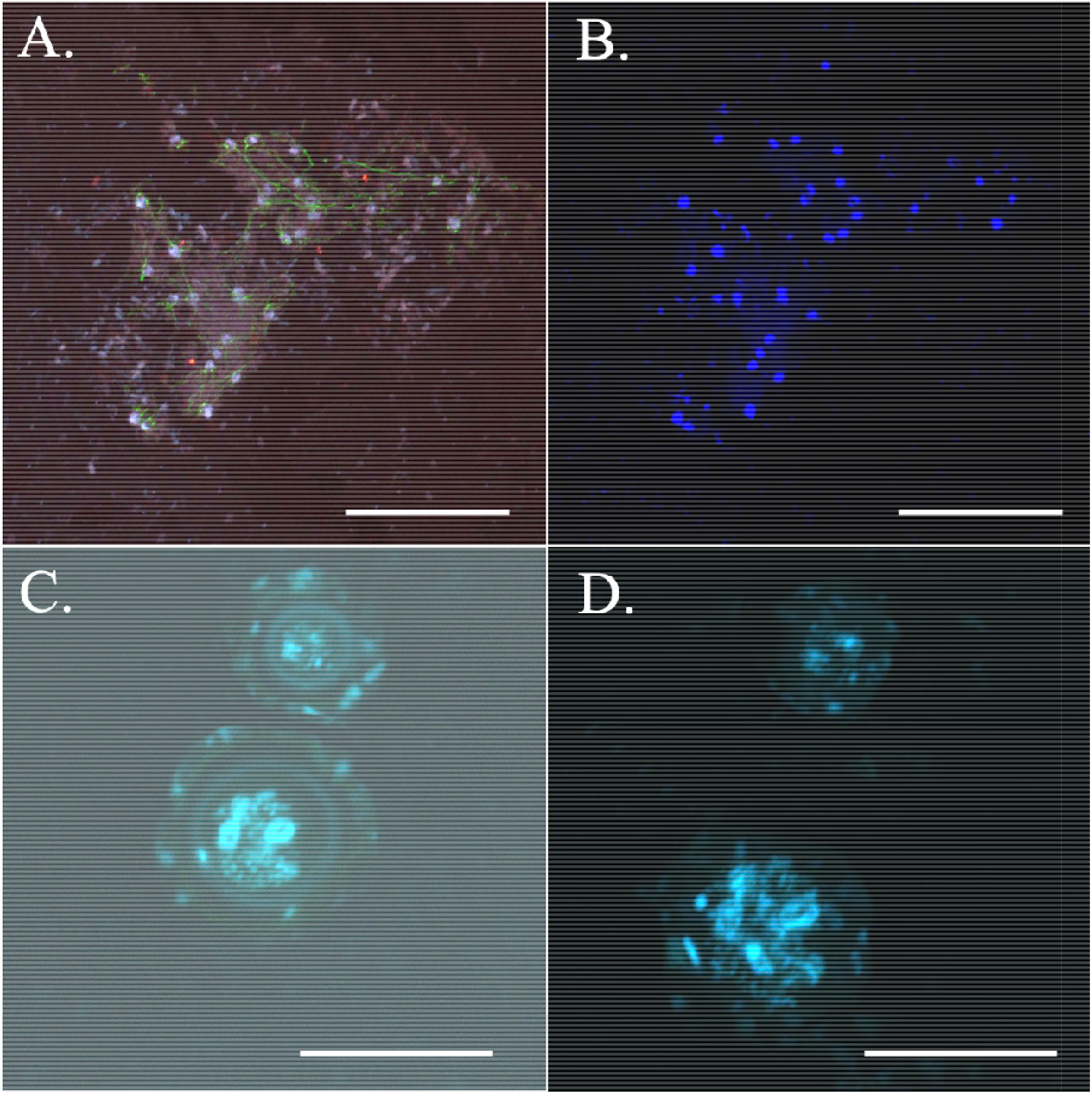
Nuclear distribution in the reticulum and cysts of *Leucomyxa quisqueyensis* sp. nov. (A) Composite maximum-intensity projection of a reticulopodial network showing α-tubulin immunolabeling (green), plasma membrane staining (red), and DAPI staining (blue). (B) DAPI channel of the same field showing the distribution of nuclei. (C) Single optical section from a confocal z-stack of two cysts. (D) Maximum-intensity projection of the same z-stack. Two DAPI-positive bodies consistent with nuclei are distinguishable in the upper cyst and at least three in the lower cyst. Scale bars are 100 µm in A–B and 10 µm in C–D.

**Fig. S2.**
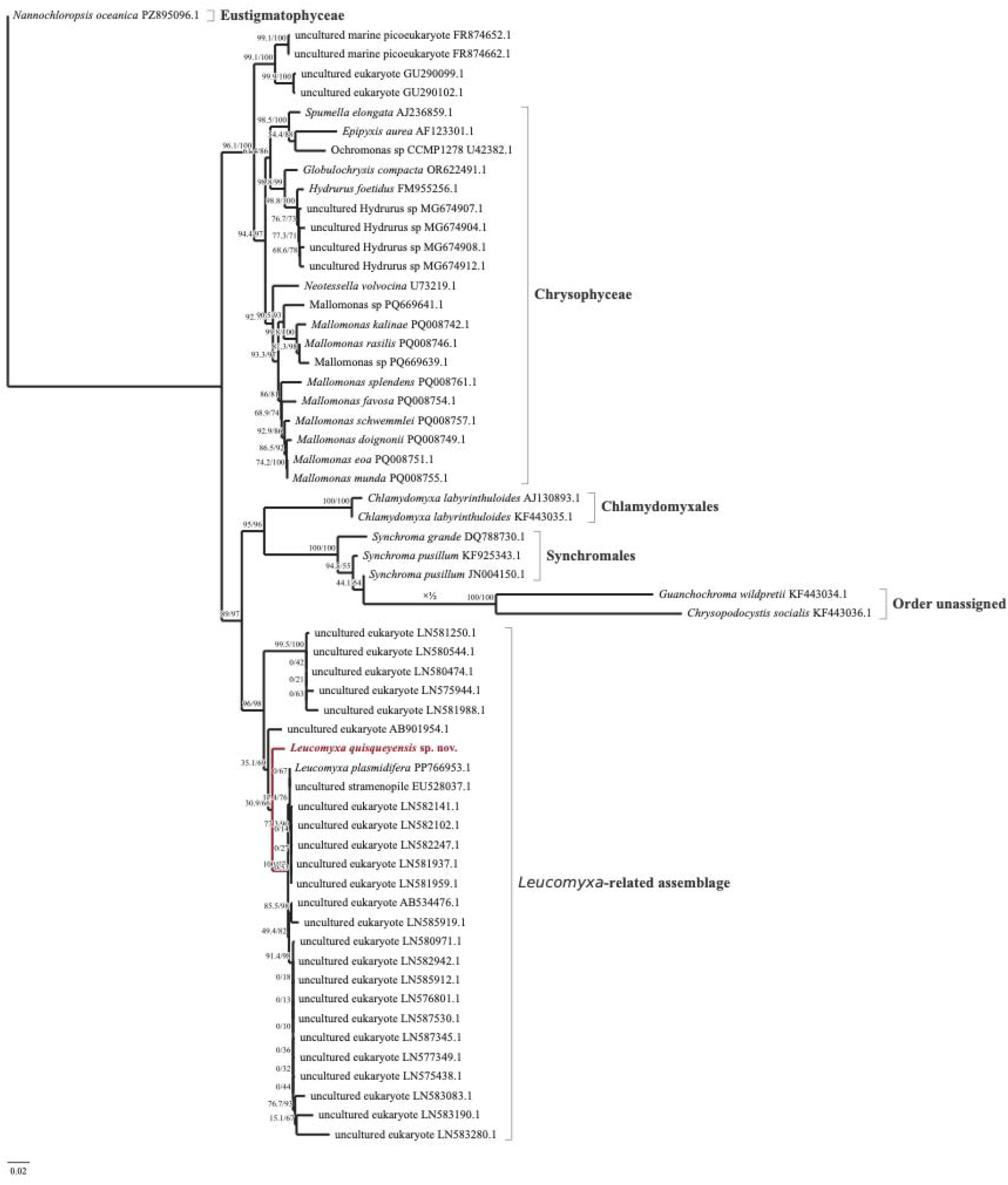
Expanded SSU rRNA phylogeny including environmental sequences. The maximum-likelihood tree was inferred from 59 sequences and 1,679 aligned nucleotide positions. It shows the placement of *Leucomyxa quisqueyensis* sp. nov. within an assemblage containing *Leucomyxa plasmidifera* and environmental sequences. The tree was rooted for display with *Nannochloropsis oceanica*. Branch labels show SH-aLRT and ultrafast bootstrap support, in that order. The scale bar represents 0.02 substitutions per nucleotide site.

**Fig. S3.**
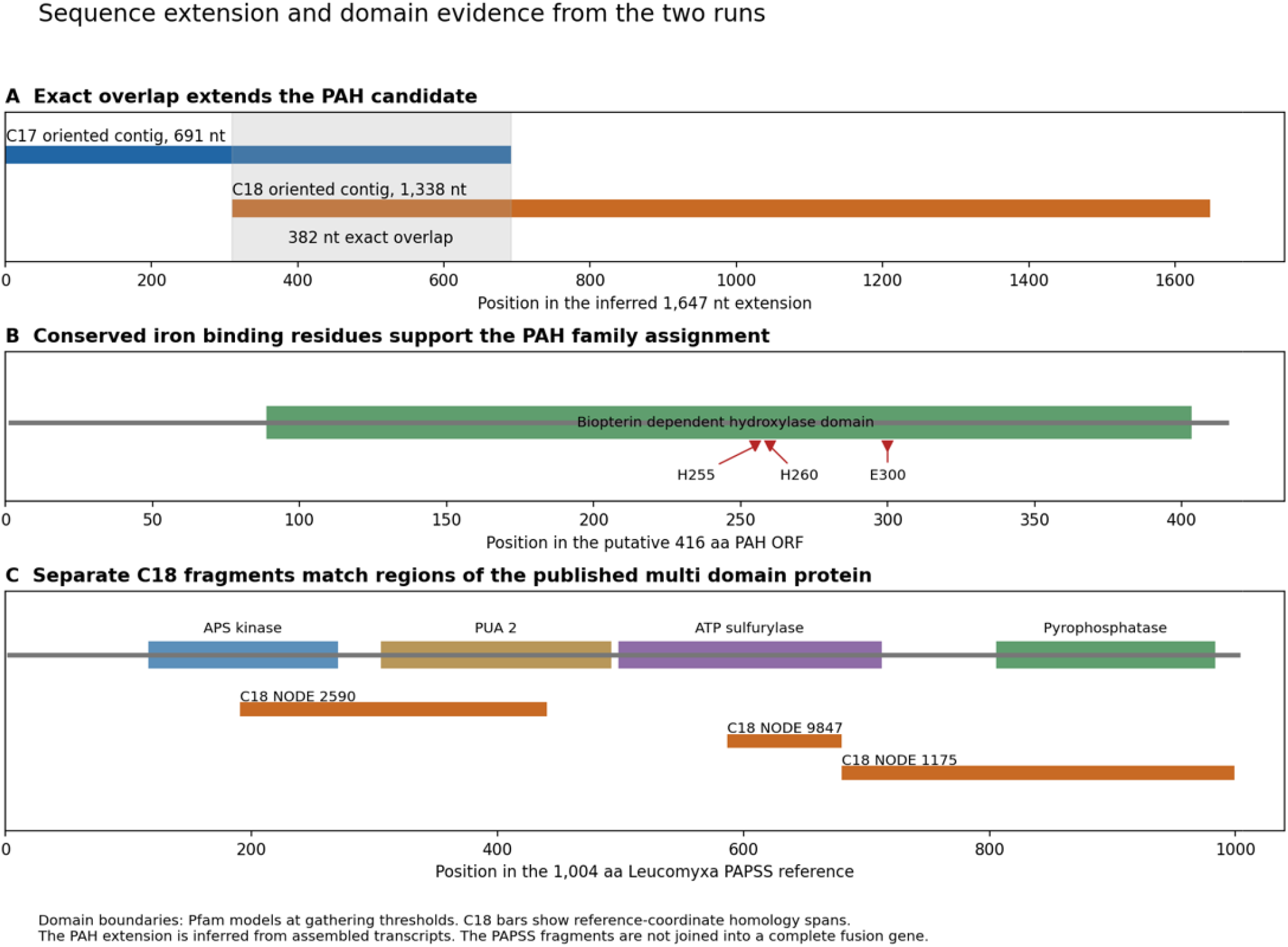
Sequence and domain evidence for PAH and PAPSS candidates in *Leucomyxa quisqueyensis* sp. nov. (A) An exact 382-nucleotide overlap between the oriented C17 and C18 contigs yielded an inferred 1,647-nucleotide sequence for the phenylalanine hydroxylase (PAH) candidate. (B) Biopterin-dependent hydroxylase domain and conserved iron-binding residues H255, H260, and E300 in the predicted 416-amino-acid PAH candidate. (C) Alignment spans of three separate C18 contigs along the 1,004-amino-acid *Leucomyxa plasmidifera* PAPSS reference protein. Colored blocks mark reference domains identified using Pfam gathering thresholds. Whether the three regions occur within a single transcript in *L. quisqueyensis* remains unresolved.

## Supplementary tables

**Table S1. Pairwise SSU rRNA sequence divergence.** For each pair, the table reports the number of comparable nucleotide positions, the number of differences, and the percentage of uncorrected divergence. Pairwise comparisons use positions with unambiguous nucleotides in both sequences. Common-site comparisons use the same 1,531 positions with unambiguous nucleotides in all five focal sequences.

**Table S2. Nuclear protein markers, taxon sampling, and individual marker-tree agreement.** (S2A) The 124 retained markers, their coordinates in the concatenated alignment, the C18 predicted protein identifiers, and the status of the C17–C18 sequence overlap. (S2B) The 38 taxa, sequence sources, and alignment occupancy. (S2C) Alignment length, reference sequence count, and whether each individual maximum-likelihood marker tree recovered *Leucomyxa quisqueyensis* sp. nov. and *Leucomyxa plasmidifera* as an exclusive pair.

**Table S3. Pairwise amino-acid divergence among the four focal taxa.** All six comparisons use the same 25,619 positions containing one of the 20 standard amino acids in all four sequences. For each pair, the table reports the number of comparable positions, the number of differences, and the percentage of uncorrected divergence.

**Table S4. Transcriptome matches to 94 published *Leucomyxa plasmidifera* protein queries.** For each query from table S5 of Barcytė et al. (2024), the table lists the functional category and detection status in C17 and C18. Where a match was detected, it gives the candidate transcript, amino-acid identity, aligned length, and query coverage.

**Table S5. Transcriptome matches to 125 protein queries derived from *Leucomyxa plasmidifera* organelle-associated reference sequences.** The queries comprise 47 proteins translated from plastid genome coding sequences (PP760456.1), 49 from mitochondrial genome coding sequences (PP760457.1), and 29 from putative mitochondrial plasmid coding sequences (PP760458.1). The table lists the reference gene and product and, where a match was detected in C17 or C18, the candidate transcript, amino-acid identity, and aligned length.

## Supplementary video

**Video S1. Reticulopodial movement and reorganization in *Leucomyxa quisqueyensis* sp. nov.** Phase-contrast time-lapse sequences show branch extension and retraction, changes in the spaces between branches, and movement of reticula across the substrate. The video includes several fields of view and is played at 10× speed. https://www.youtube.com/watch?v=JZUS_7BN1lE

